# Enterobactin and Salmochelin S4 inhibit the growth of *Staphylococcus aureus*

**DOI:** 10.1101/2024.06.30.601405

**Authors:** Yaacov Davidov, Noa Tejman-Yarden, Ari Robinson, Galia Rahav, Israel Nissan

**Author notes:** **Corresponding author:** Israel Nissan, PhD.

## Abstract

There is increasing demand for novel antimicrobial agents to tackle the antimicrobial resistance crisis. Here we report that two *Enterobacteriaceae*-produced siderophores, enterobactin and salmochelin S4, inhibit the growth of *Staphylococcus aureus* isolates, including methicillin-resistance *S. aureus* (MRSA) clinical isolates. The MIC_50_ for different *S. aureus* isolates were 2-5 μM for salmochelin S4 and 5-10 μM for enterobactin. This inhibitory activity was partially repressed by adding Fe^+3^. These siderophores also inhibited the growth of *Enterococcus* strains, including vancomycin-resistant enterococci (VRE) clinical isolates, though less effectively than for *S. aureus*. The growth of various Gram-negative bacteria was barely affected by these siderophores. These results shed new light on the role of enterobactin and salmochelin in bacterial physiology and ecology and have potential for the development of novel strategies to combat the rapid rise of multidrug-resistant bacteria.

## Introduction

New antimicrobial molecules, especially with innovative modes of action, are urgently needed to tackle the antimicrobial resistance crisis (Miethke et al., 2021). The World Health Organization (WHO) classified antimicrobial resistance as one of the top-10 global health threats faced by humanity (https://www.who.int/news-room/fact-sheets/detail/antimicrobial-resistance). *Staphylococcus aureus* is among the leading pathogens that accounts for the mortality rate associated with drug resistance, in particular by methicillin-resistant *S. aureus* (MRSA) strains (Antimicrobial Resistance, 2022). Vancomycin-resistant enterococci (VRE) are also Gram-positive resistant pathogens. MRSA and VRE are both ESKAPE pathogens which represent a global threat to human health and have been given high priority in efforts to develop new antibiotics (De Oliveira et al., 2020; Mancuso et al., 2021; Antimicrobial Resistance, 2022). The emergence of antibiotic-resistant strains has also been accelerated by the almost complete lack of new classes of clinically relevant antibiotics in the last few decades (Antimicrobial Resistance, 2022; Muteeb et al., 2023).

Iron is a mandatory nutrient for bacterial growth, due to its functions in several biological processes. However, its bioavailability is curtailed by the low solubility of ferric iron (Fe^+3^) at physiological pHs (Abbaspour et al., 2014). The human host contains high amounts of iron, but acquisition by pathogens is blocked by transport and storage proteins. Specifically, during infection, the host’s innate immune system starves pathogens by reducing intestinal iron absorption. This is achieved by increasing the production of ferritin and lactoferrin by neutrophils at the sites of infection, and by producing siderocalins (Cassat and Skaar, 2013; Nairz et al., 2014; Marchetti et al., 2020; Ullah and Lang, 2023). However, microorganisms overcome this problem by developing highly efficient uptake systems for using the iron present in the host through low-molecular weight organic chelators (150 to 2000 Da) called siderophores. These metabolites are synthesized by bacteria and released into the environment, where they chelate iron with an extremely high affinity (Johnstone and Nolan, 2015; Page, 2019; Kramer et al., 2020). Since iron uptake is essential to bacterial pathogenesis, siderophore iron uptake pathways are useful gates for antibiotic treatment using Trojan horse delivery strategies (Johnstone and Nolan, 2015; Page, 2019; Kramer et al., 2020). The tris-catecholate siderophore enterobactin is an archetype of iron acquisition in Gram-negative bacteria. It has the highest affinity for ferric iron of all natural siderophore compounds and is produced by most members of *Enterobacteriaceae* and a few other bacteria (Raymond et al., 2003). Salmochelin is a C-glucosylated enterobactin which enable it to evade the host’s defense protein lipocalin-2, an enterobactin scavenger. Salmochelins are produced by some *Salmonella, Escherichia coli* and *Klebsiella* strains (Muller et al., 2009). Salmochelin S4 is a C5,C5’ diglucosylated enterobactin and is the key compound for the production of other salmochelins (Bister et al., 2004). The ferric complex of enterobactin binds to the specific outer membrane receptor FepA, whereas ferric salmochelin binds to the IroN receptor (which is also capable of binding ferric enterobactin) (Hantke et al., 2003). The extensive research on these siderophores, their high affinity, and the ability of a variety of Gram-negative bacteria to utilize them, make them a preferred target for the conjugation of known antibiotics (sideromycins), exploiting a Trojan horse delivery strategy (Mollmann et al., 2009; Johnstone and Nolan, 2015; Page, 2019). Here we show that these two siderophores unexpectedly inhibit the growth of *S. aureus* (including MRSA clinical isolates).

## Materials and Methods

### Compounds and Bacterial Strains

Iron-free enterobactin was purchased from Sigma-Aldrich (E3910) and from EMC (Tübingen, Germany). Iron-free salmochelin S4 was purchased from EMC (Tübingen, Germany). Iron (III) chloride was purchased from Sigma-Aldrich, catalog 157740 – 100G. Cation-Adjusted Mueller-Hinton Broth (CAMHB) was purchased from BD-BBL (catalog 212322, Mueller-Hinton II Broth). Lincomycin hydrochloride was purchased from bioWORLD, Linezolid was purchased from Sigma-Aldrich (PZ0014).

The bacterial strains were kindly provided by the Clinical Microbiology Laboratory at Sheba Medical Center.

### Antibacterial activity

Antibacterial activity was determined using the broth microdilution method. The inhibitory effect was measured using broth microdilution according to Clinical and Laboratory Standards Institute (CLSI) guidelines (CLSI, 2022), in a 96-well microtiter plate, with CAMHB, and the Tecan GENios plate reader at optical density (OD) at 590nm for 18-20 h, at 37±1°C. The final bacterial inoculum was 5 × 10^5^ colony-forming units ml^-1^. MIC_50_ was defined as the lowest concentration that inhibited the bacterial growth to 50% of the final control OD. Enterobactin and salmochelin were dissolved in 100% dimethylsulfoxide (DMSO) in a stock solution of 10 mM and further diluted in double-distilled water (DDW) or directly using CAMHB to the final concentrations.

## Results

Enterobactin and salmochelin S4 effectively inhibited the growth of *S. aureus* ATCC 25923 in a dose-dependent manner (Figure 1). All the *S. aureus* strains were inhibited by these two siderophores with MIC_50_ of 2-5 μM (2-5 μg/ml) for salmochelin S4 and 5-10 μM (3.3-6.7 μg/ml) for enterobactin (table 1). Salmochelin S4 was two- to four-fold more potent than enterobactin. The growth inhibition by the two siderophores in rich media (CAMHB) at 37°C was detected after 4-5 hours of incubation and the effect was maintained for about 20 hours (Figure 1). Interestingly, a low concentration of salmochelin S4 (≤ 1.25 μM for strain ATCC 25923) or enterobactin (≤ 2.5 μM for strain ATCC 25923) enhanced the growth of the bacteria. The combination of salmochelin S4 and enterobactin displayed enhanced activity against *S. aureus* at concentrations as low as 0.5 μM for salmochelin S4 and 4 μM enterobactin as depicted in Figure 2.

**Figure 1.**
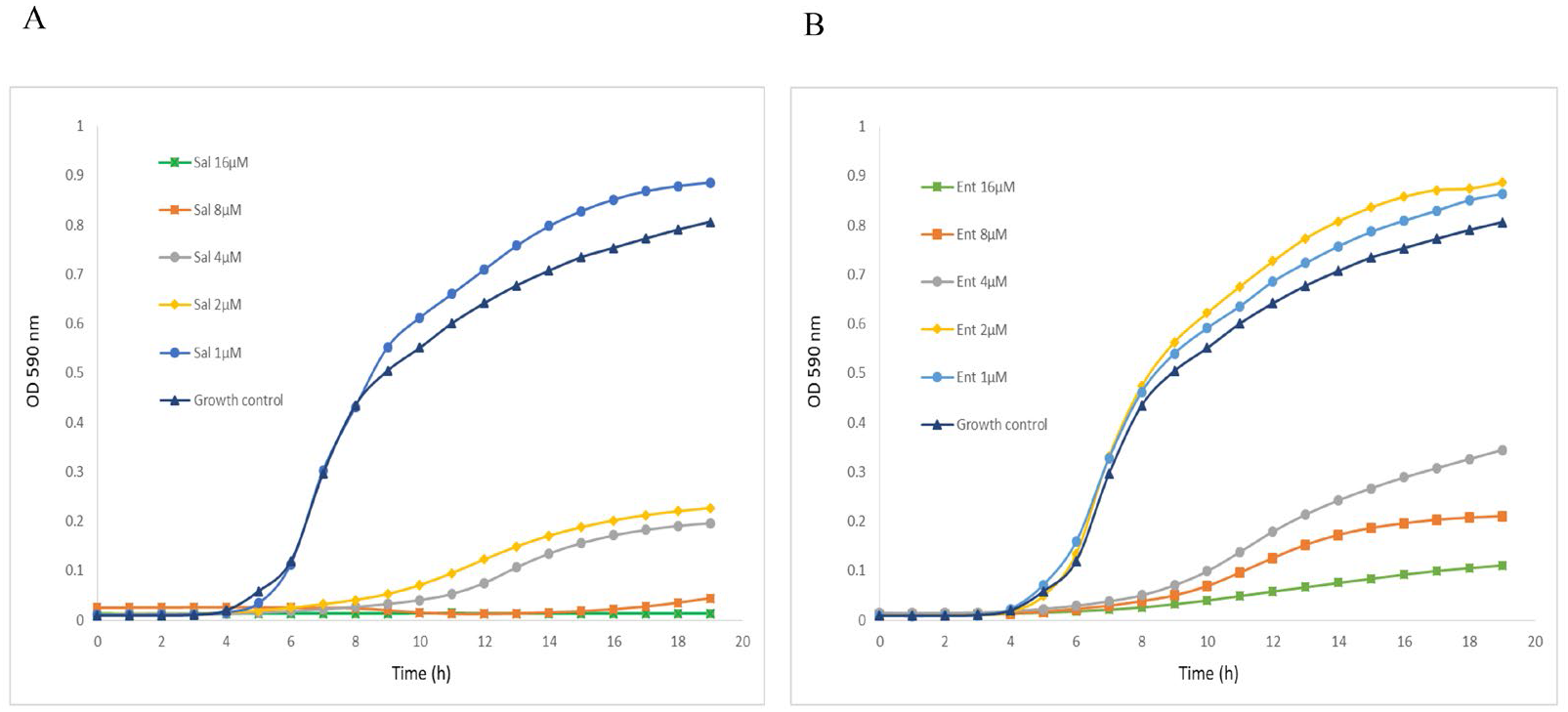
Inhibitory activity of salmochelin S4 (A) and enterobactin (B) on the growth of *S. aureus* strain ATCC 25923. Sal – salmochelin S4, Ent - enterobactin, Growth control - without siderophore.

**Table. 1:** Inhibitory effects of salmochein S4 and enterobactin on different strains of *S. aureus*.

| *S. aureus* strain | Salmochelin S4 MIC<sub>50</sub> |  | Enterobactin MIC<sub>50</sub> |  |
| --- | --- | --- | --- | --- |
|  | μM * | μg/ml | μM * | μg/ml |
| ATCC 25923 | 2.0 ± 0.4, n=7 | 2.0 | 6.0 ± 1.8, n=7 | 4.0 |
| USA300 MRSA 742 | 2.9 ± 1.2, n=9 | 2.9 | 8.4 ± 1.8, n=7 | 5.6 |
| MRSA 196 | 2.5 | 2.5 | 5.0 | 3.3 |
| USA300 MSSA 197 | 2.5 | 2.5 | 10.0 | 6.7 |
| USA300 MRSA 198 | 5.0 | 5.0 | 10.0 | 6.7 |
| MRSA 255 | 5.0 | 5.0 | > 5.0 | > 3.3 |
| MSSA 256 | 5.0 | 5.0 | > 5.0 | > 3.3 |
| ATCC 29213 | ND |  | 10 | 6.7 |

\* Main ± standard deviation. n = the number of independent experiments. ND = Not Determined. The non-ATCC strains are clinical isolates.

**Figure 2.**
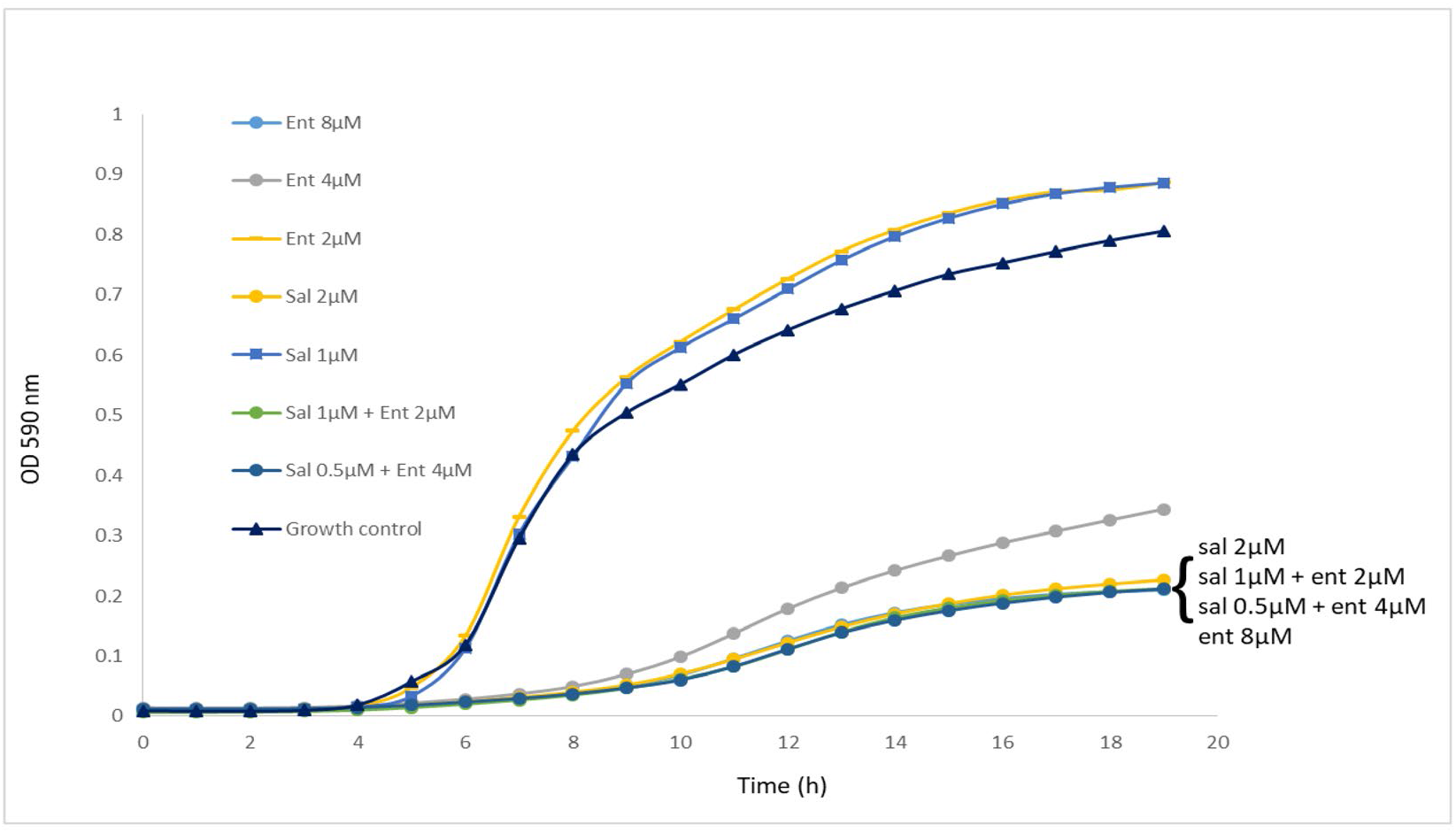
The combination of salmochelin S4 and enterobactin enhanced the inhibition of *S. aureus* strain ATCC 25923 growth. Sal - salmochelin S4, Ent - enterobactin, Growth control - without siderophore.

Next, combinations of siderophores and several antibiotics were tested. Lincomycin showed an additive effect to salmochelin S4 and enterobactin, whereas linezolid had an antagonistic effect to salmochelin S4 (Supplementary Figure 1S). These effects were also found for MRSA USA300 strain 742 (data not shown).

The inhibitory activity of enterobactin and salmochelin S4 was further tested on other Gram-positive and Gram-negative bacteria. *Enterococcus* strains, including vancomycin-resistant enterococci (VRE) clinical isolates were inhibited though at higher concentrations than for *S. aureus* (Supplementary Figure 2S). Gram-negative bacteria were not affected (*Klebsiella pneumonia, Acinetobacter baumannii*) or only slightly affected (*Escherichia coli, Pseudomonas aeruginosa*) at siderophore concentrations of up to 20 μM (Supplementary Figure 3S).

We also tested siderophore activity with addition of different concentration of Fe^+3^. The addition of Fe^+3^ reduced the growth inhibition effect of the siderophores (Figure 3). For example, the addition of 20μM Fe^+3^ to 5 μM of salmochelin S4 significantly reduced its growth inhibitory effect, whereas 10μM Fe^+3^ only had a slight effect (Figure 3A). The addition of 10-80μM Fe^+3^ without the siderophore slightly enhanced the growth of the bacteria (not shown). Co-administration of an iso-molar concentration of Fe^+3^ and the siderophores generally only had a slight effect on the inhibitory activity (Figure 3B, C).

**Figure 3.**
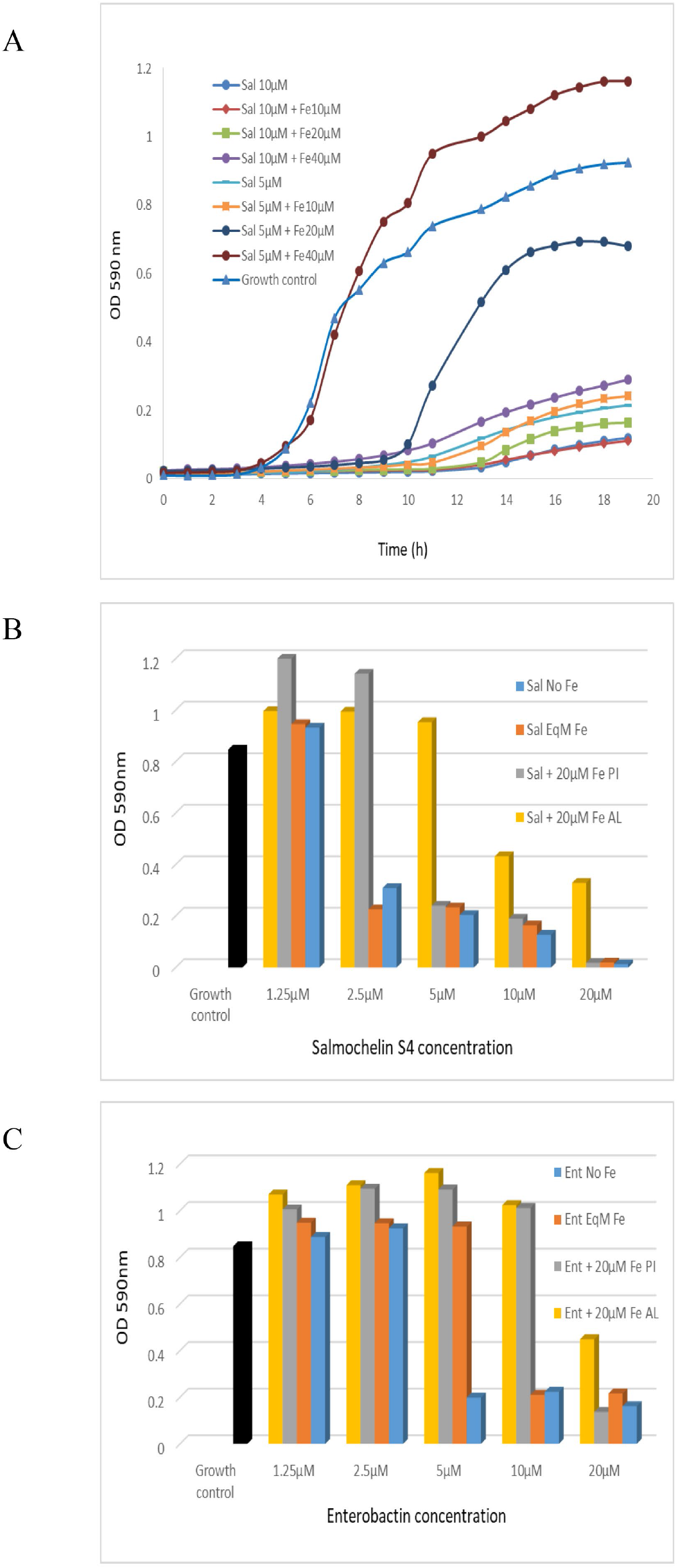
The inhibitory effect of salmochelin S4 and enterobactin with the addition of Fe^+3^ on the growth of *S. aureus* strain ATCC 25923. (A) Growth curve with the addition of 5 or 10 μM salmochelin S4 and 10, 20 or 40 μM of Fe^+3^. OD after 18 hours’ incubation with salmochelin S4 (B) or enterobactin (C). Growth control – without siderophore. Sal – salmochelin S4, Ent - enterobactin, Sal/Ent No Fe - siderophore without addition of Fe^+3^, Sal/Ent EqM Fe - Iso-molar concentration of the siderophore and Fe^+3^, Fe PI – 20μM Fe^+3^ pre-incubated with the siderophores for 30 minutes, Fe AL - 20μM Fe^+3^ added after pre-incubation of the siderophores with the bacteria for 30 minutes.

We then tested siderophore activity with addition of Fe^+3^ at different stages. The addition of 20μM Fe^+3^ when pre-incubated for 30 minutes with different concentrations of siderophores only had a slight effect on high concentrations of the siderophores (salmochelin ≥ 5 μM, enterobactin = 20 μM, for strain ATCC 25923), but a significant effect on lower siderophore concentrations (Figure 3B, C). When the bacteria were first incubated for 30 minutes with the siderophores and 20 μM Fe^+3^ were added later, the rescue effect of the iron was higher; i.e., there was less inhibition of the siderophore (Figure 3B, C). Similar results were observed for MRSA USA300 strain 742 (not shown).

## Discussion

The results demonstrate that both enterobactin and salmochelin effectively inhibited the growth of *S. aureus*, including methicillin-resistant *S. aureus* (MRSA) isolates. Salmochelin S4 exhibited greater potency than enterobactin, and their combination elicited enhanced activity. By contrast, low concentrations of the siderophores enhanced bacterial growth. Extensive research on enterobactin and salmochelin over many years has contributed to a better understanding of their importance within the bacteria that produce them, and for other organisms including *Eukarya* (Raymond et al., 2003; Muller et al., 2009; Qi and Han, 2018). The findings here contribute to furthering this field.

Several publications have described the effects of enterobactin and salmochelin on the growth of *S. aureus*. Although some of these studies have suggested that enterobactin promotes growth (Maskell, 1980; Sebulsky et al., 2000; Sebulsky and Heinrichs, 2001), or that growth is promoted by both enterobactin and salmochelin (Beasley et al., 2011), a recent study found slight growth inhibition by enterobactin for some of the *S. aureus* strains examined (Uranga et al., 2020). These inconsistencies in the impact on growth may be due to differences in media, iron availability, siderophore concentrations and other experimental conditions. Our findings make it clear that the concentrations of siderophores can either stimulate or suppress the growth of *S. aureus*.

The current data innovate by showing very effective growth inhibition for the first time of various *S. aureus* isolates including MRSA, in particular by salmochelin S4. Nolan and colleagues used conjugations of enterobactin and Salmochelin S4 to enhance activity and selectivity of β–lactam antibiotics against the pathogens that produce these siderophores. They used co-cultures with *S. aureus* (ATCC 25923) to demonstrate the selectivity and the negligible effect of the conjugations on non-producers of these siderophores (Zheng and Nolan, 2014; Sargun et al., 2021a; Sargun et al., 2021b), thus also demonstrating the relatively limited capability of *S. aureus* to absorb these siderophores (although transport through the Sst system is possible; see below).

*S. aureus*, like many other bacteria, can activate a variety of mechanisms for iron acquisition that include the secretion of endogenous siderophores, and the ability to use siderophores produced by other bacteria (xenosiderophores) (Beasley et al., 2011; Sheldon and Heinrichs, 2012; Marchetti et al., 2020; van Dijk et al., 2022). Catechol-type xenosiderophores such as salmochelin and enterobactin can be transported into the *S. aureus* cell through the Sst system (Beasley et al., 2011). The affinity of the substrate binding protein SstD for the ferric enterobactin and ferric salmochelin were found to have a *Kd* of 0.29 and 0.35 μM, respectively. These affinities are orders of magnitude lower than the affinities of the endogenous *S. aureus* siderophores to their transporters; e.g., Hts and Sir (Grigg et al., 2010a; Grigg et al., 2010b; Beasley et al., 2011). This probably reflects a sacrifice in ligand affinity in the name of greater ligand diversity (Beasley et al., 2011; Marchetti et al., 2020). Transport through the Sst system may explain the growth promotion observed when low concentrations of the siderophores are used. CAMHB, the growth medium used here, is not controlled for iron concentration. However, according to Hackel et al., (2019) the medium we used (BD-BBL, catalog number: 212322) contains 4.3μM (0.24 μg/ml) of Fe^+3^ (Hackel et al., 2019). Our results that the addition of Fe^+3^ partially represses enterobactin and salmochelin growth inhibition suggest that iron depletion is involved in the inhibition process. This depletion may be the result of a combination of the high affinity of these siderophores to iron, along with the relatively low affinity of the ferric siderophores to the SstD binding protein, and the relatively low capacity of this system to import or utilize ferric xenosiderophores, thus curtailing the availability of iron to other more effective iron acquisition systems. However, the result that Fe^+3^ addition only reduced but did not eliminate the inhibition effect, and the much stronger effect of salmochelin S4 as compared to enterobactin, may suggest that iron depletion only partially explains the inhibitory modes of action of these siderophores, and salmochelin in particular.

The potential to use iron chelators in combination with existing antibiotics was recently highlighted (Vinuesa and McConnell, 2021). The current findings provide another example of possible combinations. However, as demonstrated here, each combination should be verified independently for its efficiency.

Overall, the findings here show that enterobactin and salmochelin can act as potent inhibitors that suppress the growth of other bacterial species, thus highlighting the dual impact of these iron chelating compounds and shedding light on a novel facet of their role in bacterial physiology and ecology. Mounting evidence suggests that siderophores possess other roles beyond iron acquisition, including antibiotic activity, and can serve as mediators for interactions within microbial communities (Johnstone and Nolan, 2015; Page, 2019; Tejman-Yarden et al., 2019; Kramer et al., 2020). The report of inhibitory activity of enterobactin and salmochelin presented in this study paves the way for exploring their therapeutic applications and highlights the need for further investigation into the intricate interplay between iron acquisition and antimicrobial activity as mediated by these siderophores, including the implementation of *in viv*o experiments. Many other issues such as the breadth of this antimicrobial activity, its mode/s of action, and why salmochelin is more potent than enterobaction have yet to be discovered.

## Author contribution statement

**YD**: conducting the experiments, data curation, writing – original draft, writing – review & editing. **NTJ**: writing – original draft, writing – review & editing. **AR**: writing – review & editing. **GR**: project administration, supervision, writing – review & editing. **IN**: data curation, writing – original draft, writing – review & editing.

## Acknowledgments

We would like to extend our gratitude to Dr. Gill Smolen from the Clinical Microbiology Laboratory at Sheba Medical Center, for their invaluable assistance in providing clinical isolates used in this experiment.

