## Supplementary figures for "Enterobactin and Salmochelin S4 inhibit the growth of *Staphylococcus aureus*"

**Figure 1S.** The effect of combinations of salmochelin S4 or enterobactin with lincomycin (A) and salmochelin S4 with linezolid (B) on the growth *S. aureus* strain ATCC 25923. Sal - salmochelin, Ent - enterobactin, Linco – lincomycin, Lin – linezolid, Growth control - without siderophore and antibiotic.

A.

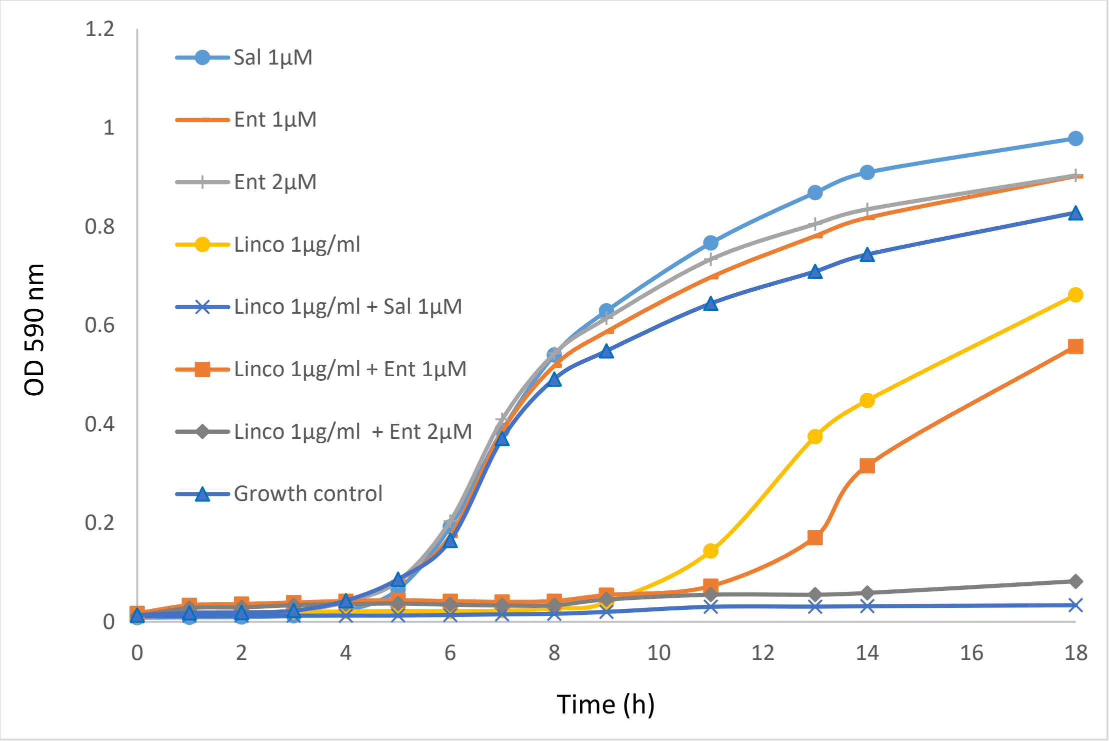


B.

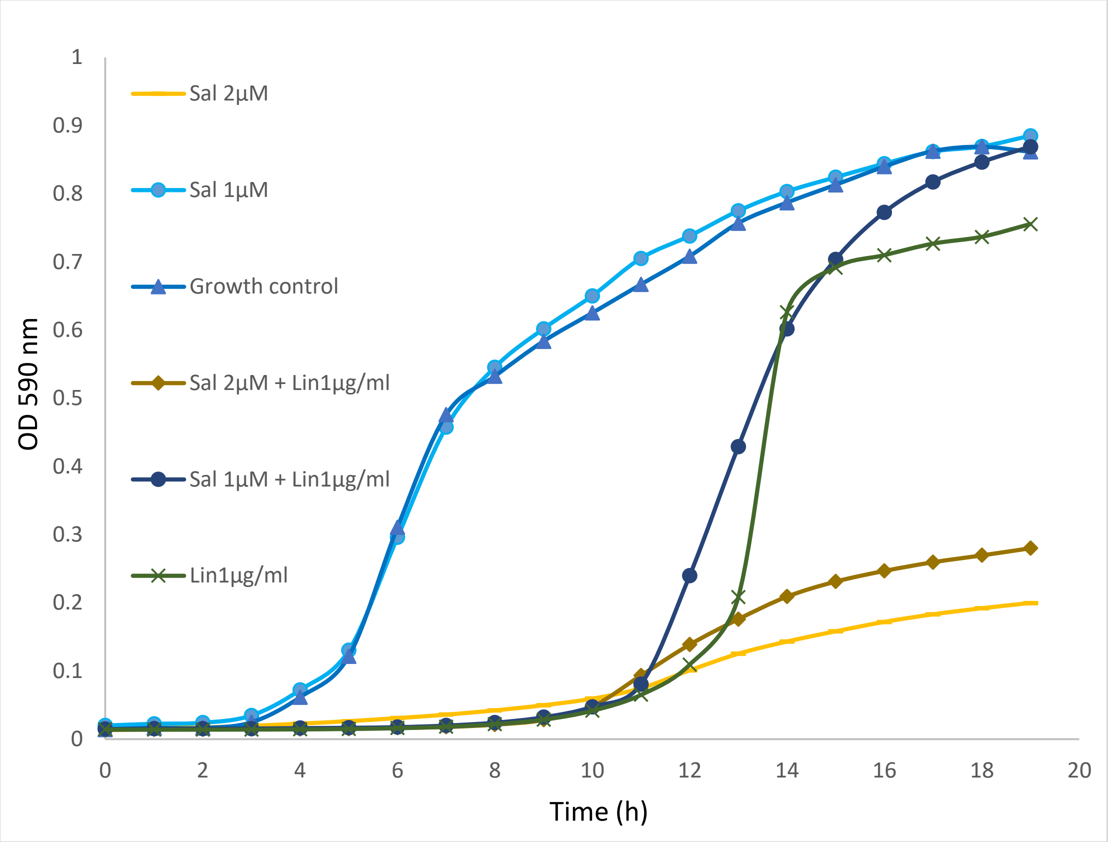


**Figure 2S**. Enterobactin (A) and salmochelin S4 (B) inhibits the growth of VRE clinical strain 195 in a dose- dependent manner. Sal - salmochelin, Ent - enterobactin, Growth control - without siderophore. Similar results were observed in other VRE clinical strains.

A.

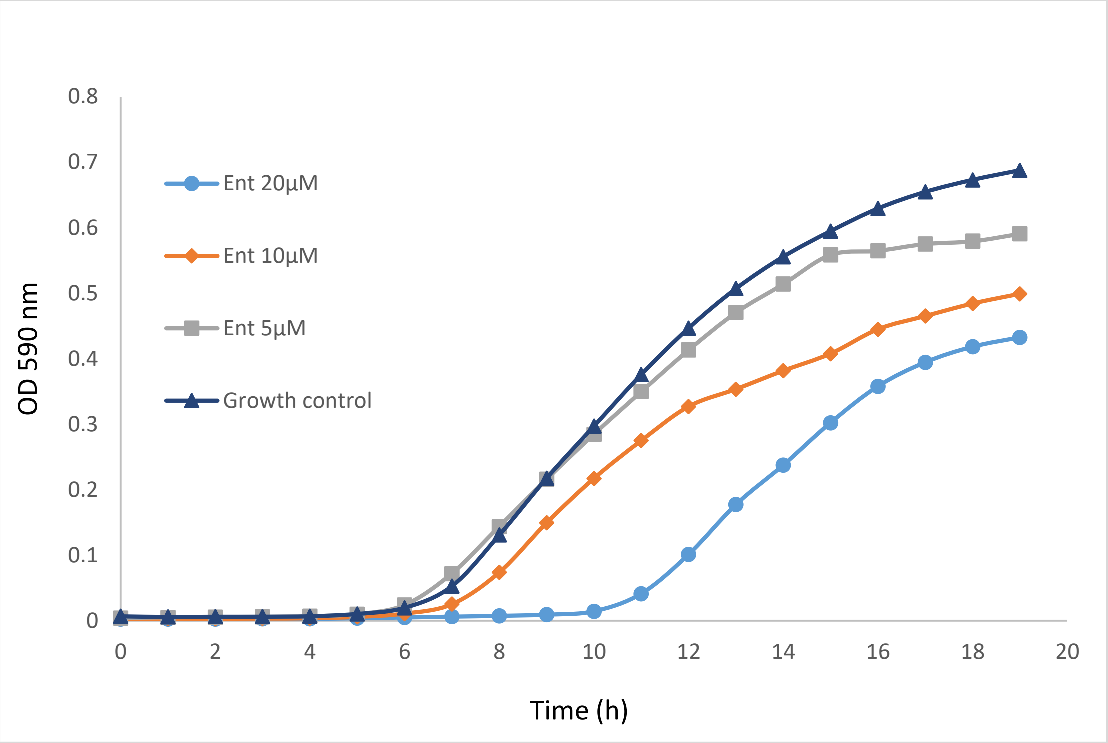


B.

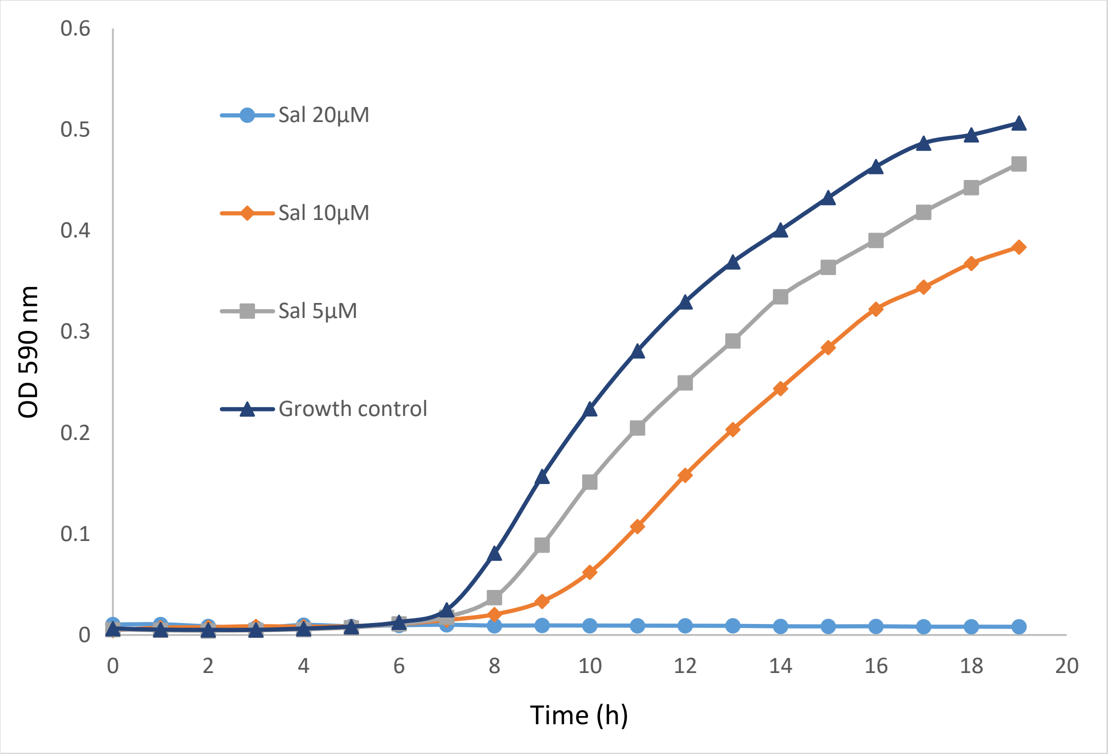


**Figure 3S.** The inhibitory activity of enterobactin and salmochelin S4 on Gram-negative bacteria. (A) *Klebsiella pneumonia* carbapenemase clinical strain (B)

*Acinetobacter baumannii* clinical strain (C) *Pseudomonas aeruginosa* ATCC 9027 (D) Uropathogenic *Escherichia coli* CFT073. Sal - salmochelin, Ent – enterobactin, Growth control - without siderophore.

A.

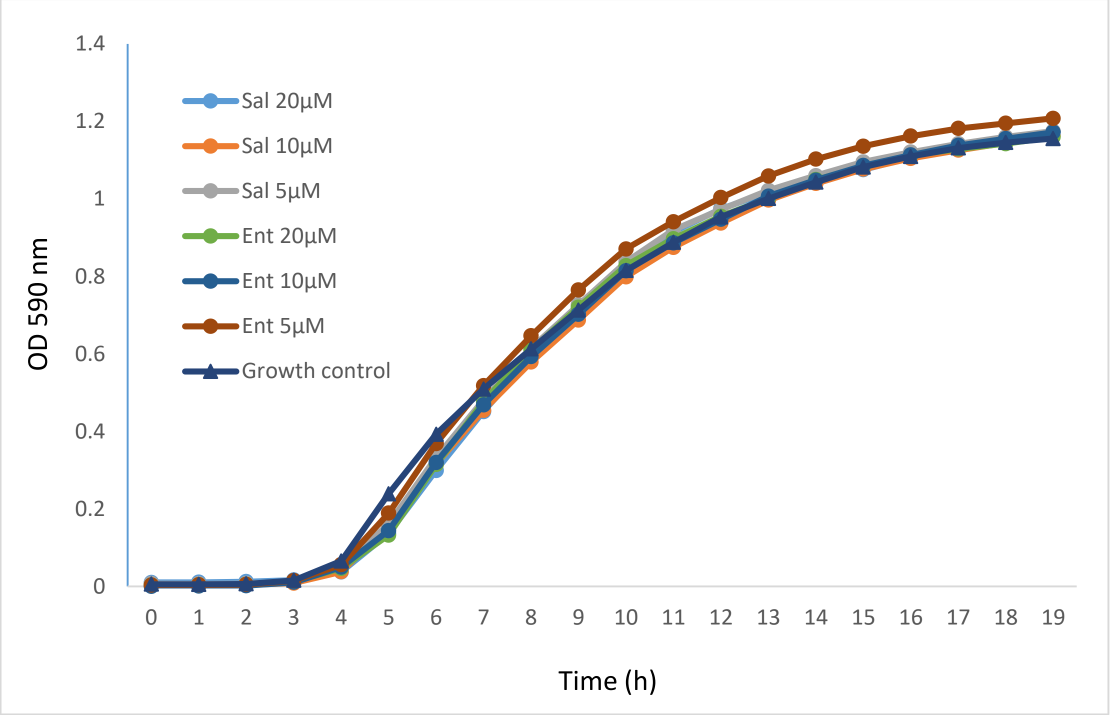


B.

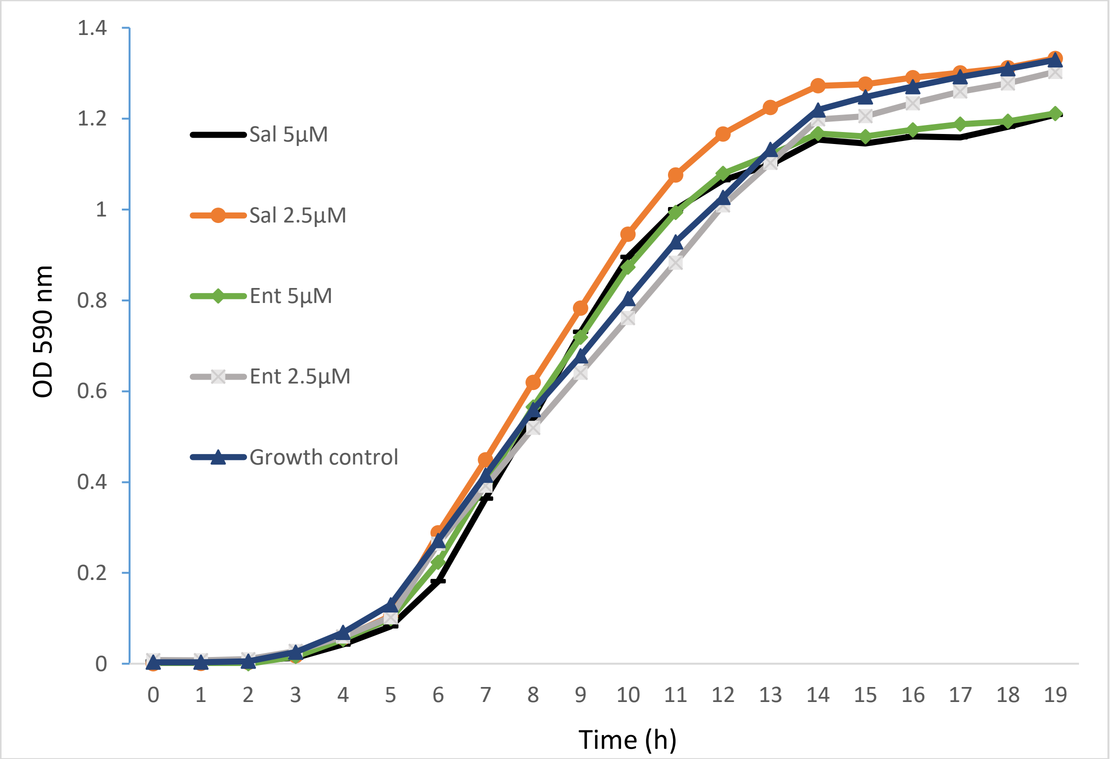

C.

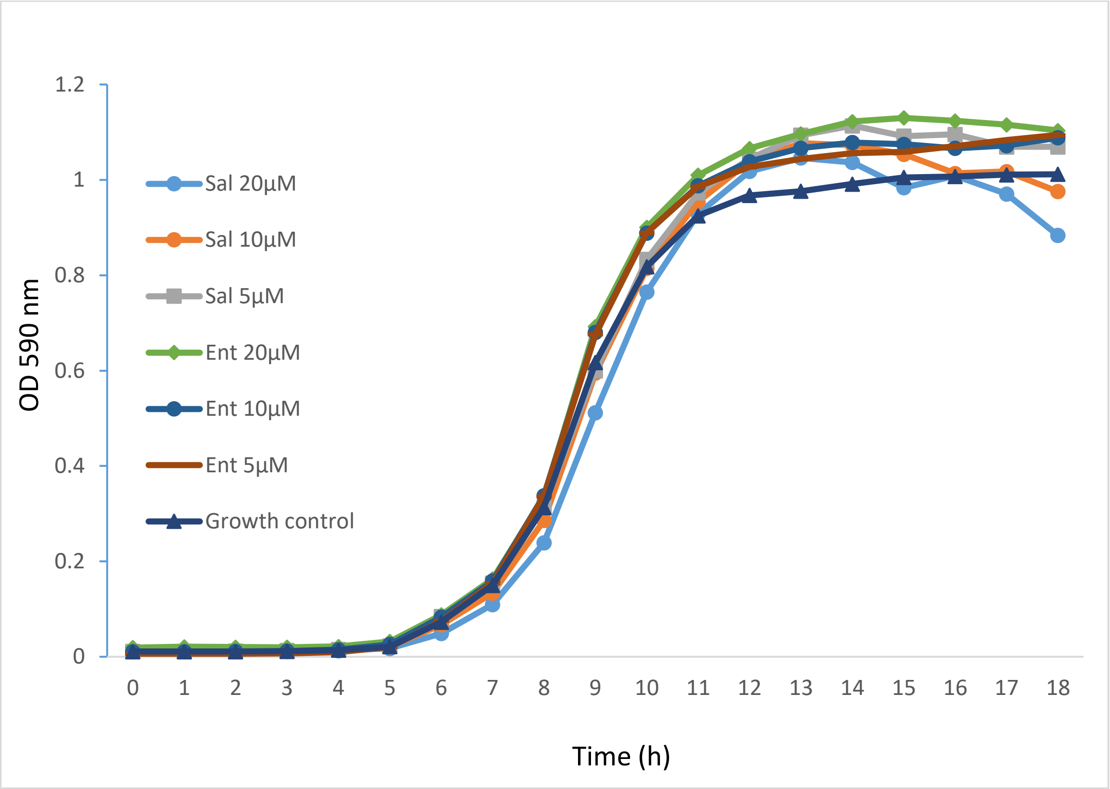
1.2
1
0.8
0.6
0.4
0.2
0
OD 590 nm
Sal 20µM
Sal 10µM
Sal 5µM
Ent 20µM
Ent 10µM
Ent 5µM
Growth control
0 1 2 3 4 5 6 7 8 9 10 11 12 13 14 15 16 17 18
Time (h)

D.

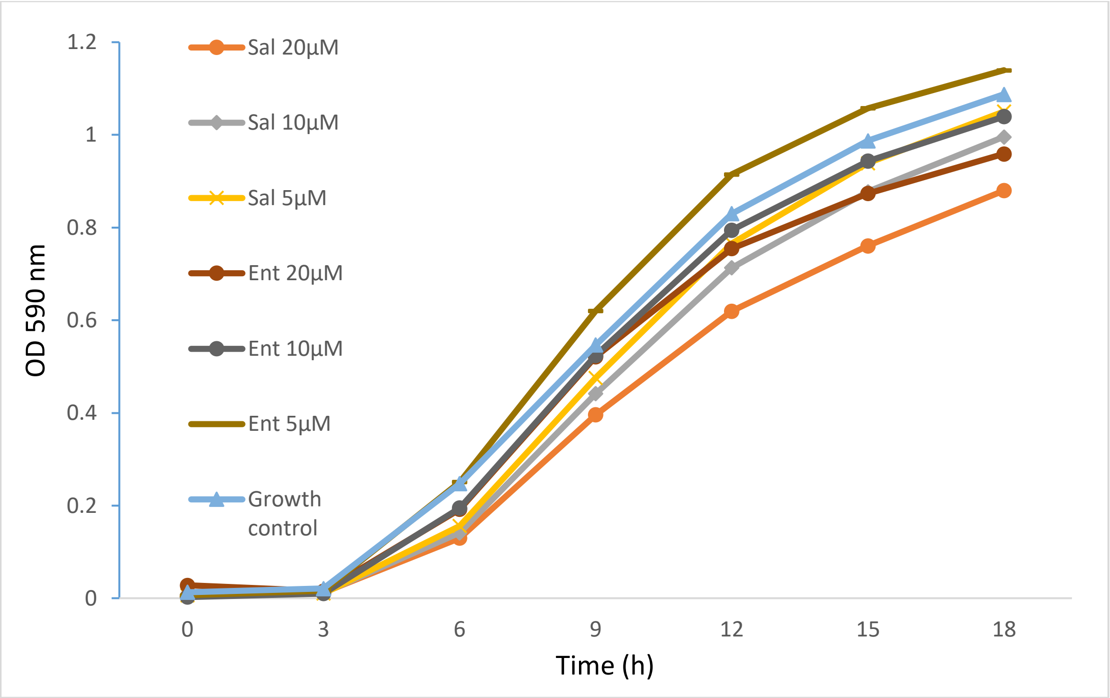
1.2
1
0.8
0.6
0.4
0.2
0
OD 590 nm
Sal 20µM
Sal 10µM
Sal 5µM
Ent 20µM
Ent 10µM
Ent 5µM
Growth control
0 3 6 9 12 15 18
Time (h)
